## Supplementary material for "Carbon dots boost dsRNA delivery in plants and increase local and systemic siRNA production"

**Table S1**. Atomic concentration table (%) for the sCD and gCD according to the XPS analysis.

| **sCD** | C1s | N1s | O1s |
| --- | --- | --- | --- |
| RSF | 0.31 | 0.50 | 0.73 |
| Corrected RSF | 6.27 | 10.18 | 15.32 |
| Atomic concentration | 74.36 | 16.32 | 9.32 |
| **gCD** | C1s | N1s | O1s |
| RSF | 0.31 | 0.50 | 0.73 |
| Corrected RSF | 6.27 | 10.18 | 15.32 |
| Atomic concentration | 73.99 | 16.70 | 9.32 |

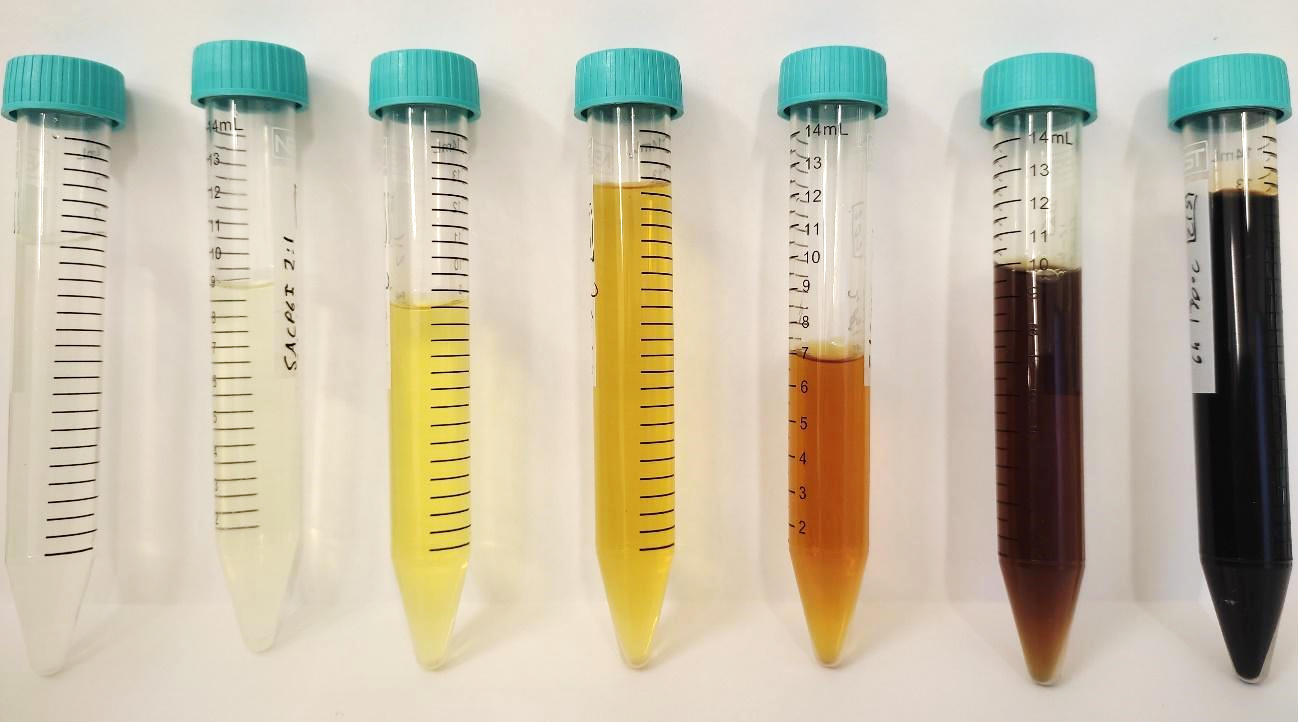

**Supp. Fig. 1**. Serial dilutions of saccharose sCDs. The aspect of the glucose gCDs dilutions resulted similar.

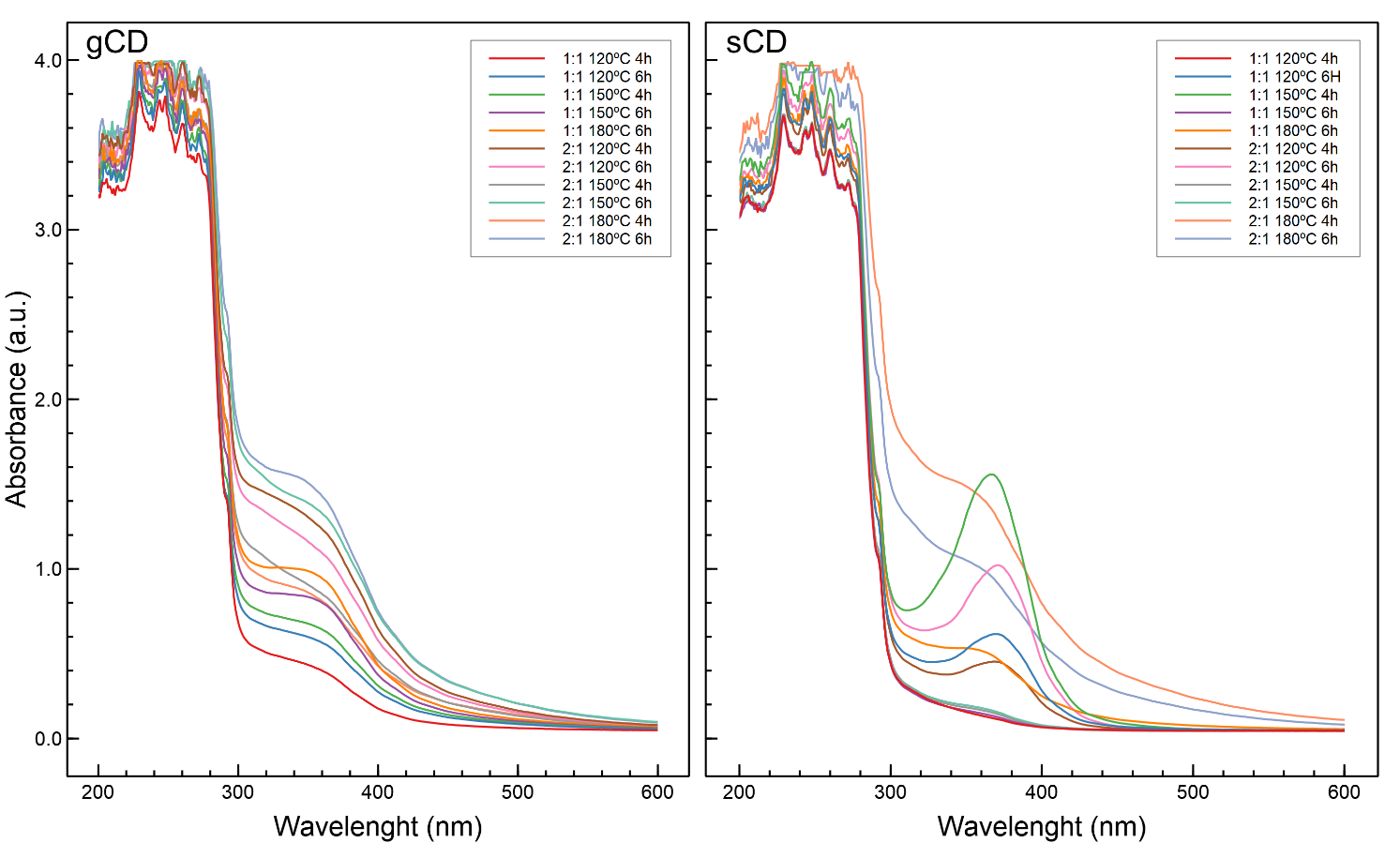

**Supp. Fig. 2**. Absorption spectra of carbon dots according to the proportion (weight: weight) between the carbon precursor and the bPEI, the temperature and time of reaction in the hydrothermal synthesis.

| A  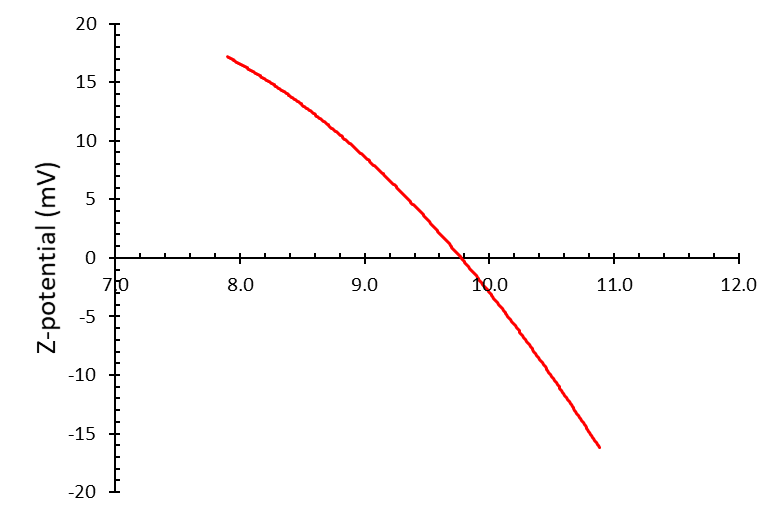 | B  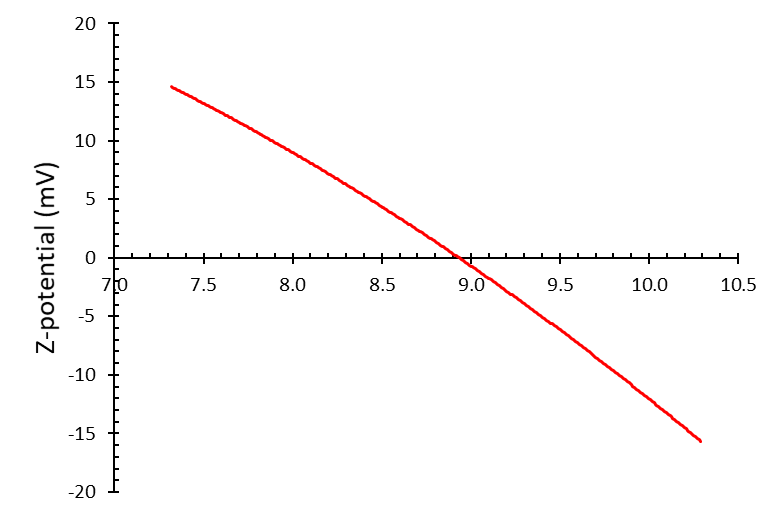 |
| --- | --- |

**Supp. Fig. 3**. Determination of isoelectric points of the carbon dots obtained from glucose (A) or saccharose (B) passivated with bPEI 2 kDa.

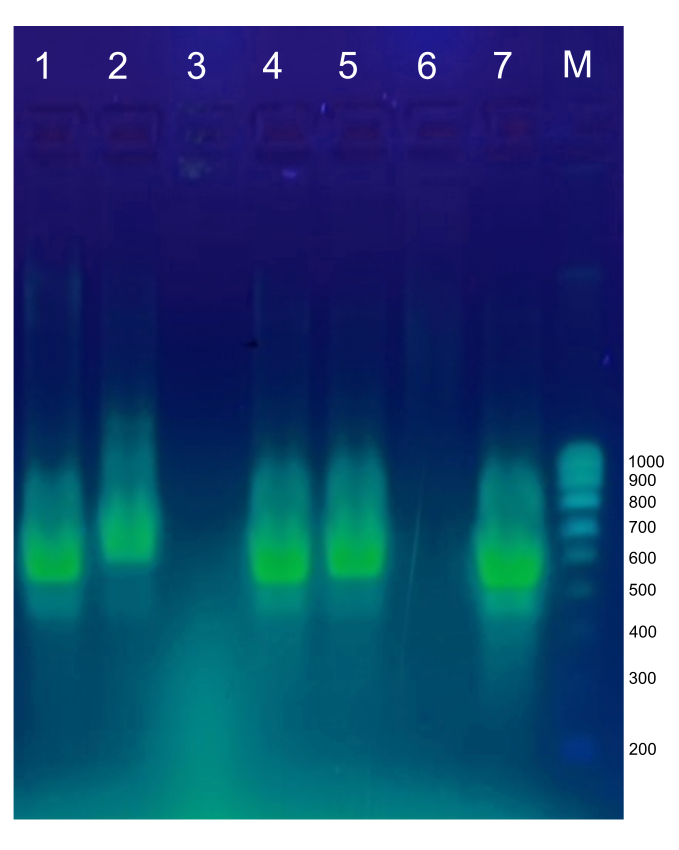

**Supp. Fig. 4**. Migration in 2% agarose gel of FITC-labeled dsRNA when naked or coated with CDs. (1) gCD:dsRNA (1:10); (2) gCD:dsRNA (1:5); (3) gCD; (4) sCD:dsRNA (1:10); (5) sCD:dsRNA (1:5); (6) sCD; (7) dsRNA; (M) Molecular weight marker: NZY Tech Ladder V.
